# Archetypes in human behavior and their brain correlates: An evolutionary trade-off approach

**DOI:** 10.1101/325803

**Authors:** Giorgia Cona, Loren Koçillari, Alessandro Palombit, Alessandra Bertoldo, Amos Maritan, Maurizio Corbetta

**Author notes:** equal contribution first author (Giorgia Cona, Loren Koçillari). equal contribution senior author (Amos Maritan, Maurizio Corbetta). **Corresponding author:** Maurizio Corbetta M.D., Department of Neuroscience, Clinica Neurologica Azienda Ospedaliera, University of Padua, Via Giustiniani 535128 Padova, Italy.

## Abstract

Organisms perform multiple tasks and in doing so face critical trade-offs. According to Pareto optimality theory, such trade-offs lead to the evolution of phenotypes that are distributed in a portion of the trait-space resembling a polytope, whose vertices represent the specialists at one of the traits (archetypes).

We applied this theory to the variability of cognitive and behavioral scores measured in 1206 individuals from the Human Connectome Project. Among all possible 300 combinations of pairs of traits, we found the best fit to Pareto optimality when individuals were plotted in the trait-space of time preferences for reward, evaluated with the Delay Discounting task. This task requires choosing either immediate smaller rewards or delayed larger rewards. Time preference for reward identified three archetypes, which accounted for variability on many cognitive, personality, and socio-economic status scores, differences in brain structure, as well as in functional connectivity between prefrontal cortex, basal ganglia, and amygdala, regions associated with reward and their regulation. There was only a weak association with genetics. In summary, time preference for reward reflects a core variable that biases human phenotypes via natural and cultural selection.

---

> *“Pleasure is the only thing worth having a theory about, ” he answered in his slow melodious voice.*
>
> *“But I am afraid I cannot claim my theory as my own. It belongs to Nature, not to me. Pleasure is Nature’s test, her sign of approval. ”*
>
> — *(Oscar Wilde, The Picture of Dorian Gray)*

According to natural selection, biological organisms maximize a specific fitness function that results in an optimal phenotype. However, when faced with complex environments, organisms need to carry out multiple tasks for optimizing survival and all of these tasks contribute to fitness. Hence organisms face a fundamental trade-off: As they cannot achieve optimal performance in all tasks, becoming specialists in one set of tasks necessarily leads to a reduction of performance in a different set of tasks. A recent paper (Shoval et al., 2012) applied a theory typically employed in engineering and economics - the Pareto Optimality - to identify evolutionary trade-offs in biological systems. Conceptualizing the space of all possible phenotypes as the morphospace, Pareto Optimality theory predicts that trade-offs among traits will lead to phenotypes (e.g., animal species) that are distributed in a small region of the morphospace. This is typically labelled Pareto Front and it is a polytope in the traits space.

The position of a given phenotype inside the Pareto front distribution is informative of its evolutionary strategy as it reflects trade-offs among traits to optimize fitness. Specifically, the vertices of the polytope are the *archetypes*, namely the phenotypes that have maximal performance in one of the tasks and minimal performance in the competing tasks. Phenotypes that fall in the middle of the polytope are *generalists*, i.e. they tend to show average performance in those tasks that define the trait space.

In the case of two competing tasks, the phenotypes fall on a line segment in the morphospace, whereas for three tasks the phenotypes fall in a triangle. Four tasks result in a tetrahedron distribution, and so on. Notably, this analysis is data-driven since it is the distribution of the data to indicate which tasks show trade-offs.

The Pareto Optimality approach has been successfully applied to datasets from animal behavior (Gallagher et al., 2013), animal morphology (Shoval et al., 2012; Szekely et al., 2015), cancer (Hart et al., 2015), bacterial gene expression (Thøgersen et al., 2013) and biological circuits (Szekely et al., 2015).

For instance, the study by Szekely et al. (2015) demonstrated that animal species when analyzed in terms of size and longevity fall on a triangular distribution, with the vertices representing the three archetypes: large animals with high longevity (e.g. whales being the archetype), small animals with high longevity (e.g. bats), and small animals with low longevity (e.g. mice). These specialists enrich on other attributes, such as fertility and predation. For instance, small animals with low longevity tend to have high fertility and to be preys while small animals with high longevity tend to have lower fertility but also low predation.

The present study tested for the first time the predictions of Pareto optimality theory to human cognition and behavior. Specifically, in a large population of subjects, the theory predicts that some individuals will excel in some tasks at the expense of other tasks (i.e. *archetypes)*, while others (i.e. *generalists)* will show average performance on most tasks. It is important to emphasize that a Pareto distribution, i.e. one showing trade-offs in tasks or behaviors, is not a given. In fact, this theory contrasts with the existence of a general factor of intelligence, or g, which has been inferred from the robust positive correlation (across subjects) detected across a large number of cognitive tasks (Spearman, 1904). While human intelligence may embrace more than sixty specific cognitive abilities, an underlying common factor (*g factor)* is common to all of them (Carroll, 1993; Colom et al., 2006), and explains large amount of variance (45-50 %) across tests scores in large samples of healthy subjects (Austin et al., 2002; Floyd et al., 2009). In contrast to Pareto optimality theory, the g factor theory predicts that different cognitive or behavioral variables will not show a trade-off, but only positive correlation across subjects.

To test the predictions of Pareto optimality theory in human cognition and behavior, we analyzed data from the Human Connectome Project (HCP) that includes a wealth of cognitive, personality, health, socio-economic status, and brain measures (Van Essen et al., 2013). Specifically, we first identified the best triangle that explained the highest fraction of variability across subjects. The vertices of such triangle define the archetypes, i.e. the specialists for that set of tasks. Next, we defined the traits that characterize (enrich) the archetypes, by taking into account not only cognitive, but also personality, and socio-demographic measures. Based on this enrichment analysis, we inferred the competing human evolutionary strategies. Finally, we identified differences among archetypes in brain structure (volume, gray matter, etc.), function (resting state functional magnetic resonance imaging, rs-fMRI, connectivity), and potential genetic influence.

## Materials and Methods

### HCP Dataset

We analyzed the public data release of the WU-Minn Human Connectome Project (HCP) consortium (van Essen et al., 2013), which released a publicly available dataset of 1206 healthy young adults, from families with both twins and non-twin siblings. The current sample was obtained from the March 2017 data release (1200 Participants; http://www.humanconnectome.org). The database consists of behavioural measures (e.g., cognitive, personality), socio-demographic measures, and high-resolution 3T MRI imaging data.

Some data are restricted due to subject privacy (e.g. which subjects are twins, which smokers etc). The HCP subjects include 168 Monozygotic twin pairs, 103 Dizygotic twin pairs. The behavioral database consists of tests that are part of the NIH Toolbox battery and of several Non-Toolbox behavioral measures (see Supplementary Material for a more detailed description of the measures).

For each subject, we also obtained the brain volumes from the Freesurfer software and analysed them by voxel-based morphometry. They consist of continuous features and are normalized with respect to the intracranial volume.

### Pareto Optimality Inference method

The Pareto Optimality analysis is based on the following assumptions:

1. Each subject is featured by a vector of continuous traits *ν*, obtained through measures of cognitive, personality, socio-demographic, and brain features in the case of the HCP dataset.
2. Each subject can perform *k* multiple tasks, which are in competition between each other. Each task is assigned a performance function P_k_, which quantifies the ability of a given subject to perform that task.
3. The performance of a given task is maximal at a given vertex (which is populated by subjects that are specialists in that task, a.k.a. *archetypes*) and the performance decreases with the Euclidean distance from the archetypes.
4. The fitness function of a given subject is an increasing function of the performance at all tasks, F(P_1_(ν)…P_k_(ν)).

The maximization of the fitness function is a multi-objective optimization problem that has as solutions sets of points enclosed in polytopes in the space of traits (Shoval et al., 2012). The points nearest each vertex of the polytopes correspond to subjects specialized in tasks with the highest performances in that task and lowest in the others. The Pareto Optimality analysis is performed in order to identify: 1) the best-shaped polytope that encloses the data points in the space of traits; and 2) its vertices, which would represent the archetypes, namely clusters of specialized individuals in features that are in trade-off with each other (Hart et al., 2015). The predictions of Pareto Optimality theory are indeed met only if both conditions are satisfied.

As compared with the classical clustering methods (k-means, Gaussian Mixture models, Latent Class Analysis), Pareto Optimality approach differs as it identifies the extreme vertices (rather than centroids) of a distribution. Clustering and Pareto analysis are indeed both able to find centroids, but in a complementary way, since the former is sensible to local density, while Pareto is sensible to the external shape of distributions, also called convex hulls (for further differences between the Pareto method and clustering see Hart et al., 2015).

The first step in our analysis was projecting for each pair of behavioral measures the 1206 participants’ data points in a two dimensional space. We considered twenty-five measures related to each cognitive and performance domain (e.g., fluid intelligence, memory, spatial orienting, self-regulation, strength, dexterity etc. (see Supplementary Materials). After removing redundant, ordinal measures and measures with too few available values, 300 combinations of pairs of cognitive and performance-related traits were tested.

As a second step, we checked for the best-shaped triangular hull among all possible combinations of traits. The statistical significance of the convex hull was achieved by using the triangularity test (the t-ratio test). The t-ratio stands for the fraction between the area of the triangular hull, performed through Sisal algorithm (Bioucas-Dias, J. M. 2009), and the area of the best convex hull that encloses the cloud of data points, typically with a higher number of vertices. The t-ratio values closer to 1 indicate a better fit of the cloud to a triangle. For each triangular-shaped distribution we tested the robustness of the triangles by comparing the t-ratio of the original distributions with the t-ratio derived from n-null distributions obtained through 1000 random permutations of the values of the data points, i.e. by leaving the same cumulative distribution function (CDF) for a given set of values. This defines the p-value as the fraction of times the null t-ratios are lower than the empirical t-ratio, and statistical significant p-values should score under 5% of times (p <.05). We performed this analysis in the space of each of the 300 combinations of traits, and in each case we found a p-value for the triangular-hull. A correction for multiple comparisons was applied to all the p-values thought the False Discovery Rate (FDR) method.

To further control the triangularity of the Pareto distribution, we measured the fraction of variance accounted for (across subjects) as a function of the number of vertices (2 to 6) of the possible polygons.

### Reliability of Pareto Front Solution

Even though the triangularity test examines the statistical significance of the obtained Pareto front solution against a null distribution through permutation tests, we also ran additional control analyses to validate our findings.

In one control analysis, we performed a split-half replication: we ran the Pareto analysis separately on two random independent sub-samples of the HCP data set (n=559 and n=560 subjects, respectively), taking into account all 300 possible combinations of pairs of traits. This was done to ensure that the Pareto Front solution obtained from Pareto Optimality Inference method was robust, i.e. significant in two separate samples.

In two additional control analyses we asked whether the obtained Pareto front solution was robust to gender and race. In one analysis, subjects were split the subjects by gender: Males (549 subjects), Females (649 subjects). In the second analysis, subjects were split by Race: Asian-Nat. Hawaiian-Other Pacific (n=67 subjects), Black or African American (n=192 subjects), White (n=883 subjects).

### Enrichment analysis of the Archetypes

According to the Pareto Optimality theory, the vertices of the triangle identify specialists that maximally (o minimally) express different traits. From this set of traits, one can infer traits/tasks that are in trade-off between each other. If Pareto theory is correct, then, other traits (i.e., enriched features) should be maximal or minimal in those individuals who are located closest to the vertices (i.e., archetypes), and should decline (or rise) as a function of the distance from that archetype.

To identify the traits that enrich the three archetypes of the optimal Pareto front solutions, we first divided the distribution in bins and then analyzed, for each trait, the change of the mean value of that trait across the bins of the polytope, normalized with respect to the mean value of the given trait for the whole distribution. For simplicity, we binned the Pareto front three times, each time starting from one of the three vertices, into n bins. To make the analysis statistically valid in terms of sample size, we constrained each bin to contain the same number of participants. This procedure was repeated systematically by varying the number of bins between 8 and 15. A higher number of bins leads to higher statistical fluctuations in the density analysis. Features could be discrete or continuous. For continuous variables, we computed the ratio among the mean value at all bins and the mean value of the entire triangle. We plotted this ratio as a function of the n-th bin. For discrete features, we first booleanized them (i.e. a value 1 was given if the participant had the given feature, 0 otherwise), then we treated them as continuous variables.

Enriched features were validated if they passed the p-value test (based on the hypergeometrical distribution (Hart et al., 2015) and corrected for FDR test), which measures the probability that the mean value of a trait is maximal/minimal in the bin closest to a given vertex. The robustness of the enrichment was assessed by performing a null-test, namely a random permutation of the values of the traits among the different bins. Features belonging to four main domains were separately analyzed: 1. Cognitive, Physical and Sensory traits (1119 subjects and 46 measures); 2. Discrete traits of Personality, Habits, socio-demographic features (1123 subjects, 40 measures); 3. Continuous traits of Personality, Habits, socio-demographic (1123 subjects, 70 continuous measures); 4. Structural brain measures (1105 subjects and 56 measures). Structural brain measures (n=56) included volume of cortical gray matter, white matter, and volume of anatomical regions in the right and left hemisphere (e.g. right and left hippocampus, thalamus, etc․.) segmented in Free Surfer. Before running the enrichment analysis, the measures were first normalized per intracranial volume.

### Resting-state Functional Connectivity analysis

To characterize differences in functional connectivity among different archetypes of significant Pareto front solution, we analyzed resting state functional connectivity (FC) from R-fMRI as available in the HCP data set.

#### Subjects

Three-hundred healthy subjects (172 F, age: 29 ± 3y) were selected from the 1200-subject release HCP dataset, considering, for each archetype, 100 subjects with minimal Euclidean distance from each archetype vertex of the Pareto distribution. This sample size was selected because it was similar to the average sample size of the binning analysis for feature enrichment.

#### Imaging Data

The HCP imaging protocol included up to 4 runs of resting state fMRI of 15 minutes of duration (TR = 720ms, isotropic voxel-size 2 mm) and structural images, made available as data packages with pre-defined processing options (Glasser et al., 2013). In this analysis, we employed minimally pre-processed fMRI time series from surface space defined and registered by means of a Multi-modal surface alignment method (MSM-All, (Robinson et al., 2014)) with minimal smoothing (surface and volume based 2mm spatial smoothing) and de-trending. Moreover, FIX-ICA (Salimi-Khorshidi et al., 2014) denoised data was employed as available from HCP public repository to reduce motion-related confounds (Marcus et al., 2011).

#### Data Processing

Available denoised rs-fMRI time-series were signal averaged based on the functional parcels defined from the Gordon-Lauman scheme (2016) for cortical regions, and a volume based segmentation (Fischl et al., 2002) for subcortical regions (Cerebellum, Putamen, Pallidum, Ventral Diencephalon, Thalamus, Caudate, Amygdala, Hippocampus, and Accumbens in each hemisphere and Brainstem). Parcellated rs-fMRI time series were Pearson cross-correlated and Fisher r-to-z transformed (z = 0.5 * ln[(1 + r)/(1r)], with r the estimated Pearson linear correlation coefficient at edge-level (Hilinka et al., 2011) to obtain for each subject and run a FC matrix across 352 brain regions (Smith et al., 2011). We discarded rs-fMRI runs that included more than 30% of motion corrupted volumes. Framewise Displacement (FD) was employed to identify the motion-corrupted volumes as it indexes bulk head movements across consecutive volumes (Power et al., 2014) from the volume realignment parameters (motion correction). Since the available rs-fMRI data were previously pre-processed with FIX-ICA denoising, we relaxed the threshold for motion-corrupted volumes to FD > 0.5 mm as compared to previous suggestions of FD > 0.15 - 0.2 mm (Power et al., 2014). After removal of motion-corrupted runs, all subjects had at least two valid fMRI runs. Correlation values in corresponding edges were finally averaged across valid runs to obtain a single FC matrix per subject. The subjects included in the sample were not found to be significantly different in terms of motion content as function of the archetype. Inter-run and inter-subject global variability was removed by normalization (Geerligs et al., 2017).

We performed a region of interest (ROI) analysis in the three groups of subjects based on a-priori hypotheses in the literature concerning the cortical and subcortical regions recruited by the tasks showing significant and replicable Pareto front solutions (see below, results section). Regarding the results for DDT-related Pareto front, we considered ROIs from recent studies and meta-analyses of the DDT and reward processes (Liu et al., 2011; Li et al., 2013; Wesley and Bickel, 2014). The selected ROIs were: Ventromedial prefrontal cortex (vmPFC), orbitofrontal gyrus (OFG), middle frontal gyrus (MFG), dorsomedial prefrontal cortex (dmPFC), dorsolateral prefrontal cortex (dlPFC), superior frontal gyrus (SFG), anterior prefrontal cortex (aPFC), anterior cingulate cortex (ACC), posterior cingulate cortex (PCC), anterior internal capsule (aIC), hippocampus (Hip), parahippocampus (Parahip), Striatum, Caudatum, Putamen, Accumbens, Globus Pallidus, Thalamus, and Amigdala. The correspondence between the literature ROIs and the 352 Gordon-Lauman defined cortical/subcortical regions/parcels was established according to a visual spatial overlap criterion at the group level. In general, we found that regions of activation tended to involve multiple adjacent parcels of the 352 Gordon-Lauman parcellation scheme. The selected ROIs overlapped with 63 parcels of the Gordon-Lauman atlas extended with subcortical regions. Therefore, the initial 352×352 FC matrices were reduced to 63×63 matrices describing the connectivity of selected regions. However, even this smaller matrix tended to oversample the region of activations. Since we did not observe significant differences in connectivity profiles among neighboring parcels near/at the focus of activation, and corresponding homologous regions, we further reduced the matrix to 18×18 showing distinct connectivity profiles.

#### Analysis and statistical comparisons

We carried out a Ward hierarchical clustering between coupled archetypes based on Euclidean distance similarity of connectivity profiles (i.e. FC rows, or columns by symmetry) similar to Nomi and Uddin (2015). This analysis consists in the hierarchical clustering of FC matrices to identify the node clustering structure of one group of subjects (e.g. those belonging to one archetype) and use this structure to reshape the FC representation of another group of subjects (those belonging to the other archetype). In this way, differential hierarchical organization between FC in different groups of subjects will be visually clarified. As we did not find any significant difference in the FC hierarchical organization among the three archetypes, the reported analysis is based on clustering of FC matrices based on all subjects across the three groups. Next, we tested for differences among groups using a 1-way Analysis of Variance (1w-ANOVA) with bootstrap sampling for statistic evaluation on pair-wise ROI FC (Fisher-transformed Pearson correlations) testing the null hypothesis of equal connectivity between the three archetypes (see Xu et al., 2014, for a similar approach). An FDR method was applied to correct for not independent multiple comparisons testing conditions. Post-hoc tests were run by means of one-tailed paired two-sample t-test with bootstrap sampling to investigate the directionality of connectivity by archetypes couples. FDR correction was again employed and restricted according to a Bonferroni strategy over the number of performed post-hoc tests.

#### Software and tools

Processing of rs-fMRI data, available as Neuroimaging Informatics Technology Initiative volumes (NIFTI) or Connectivity File Based Data (CIFTI) files was done with Connectome Workbench (Marcus et al., 2011) and CARET (Van Essen Laboratory, Washington University) as well as surface visualization and representation of relevant brain areas. Statistical comparisons and further analysis were performed in MATLAB (R2016b; MathWorks, Natick, MA).

## Results

### A Pareto Front distribution for the delay discounting task (DDT)

For each participant, we took into account 25 continuous measures of the HCP (i.e., cognitive and behavioural scores), mapping them into the multi-dimensional space of traits (i.e., morphospace).

The best triangular Pareto front solution was found in a two dimensional space that contains, for each subject, the values associated with the Area-under-the-curve (AUC) for $200 and AUC for $40,000, two measures of the delay discounting task (DDT) (Figure 1). Indeed, among all possible pairwise combinations of traits, the triangle defined by the two measures of the DDT was the only one to survive the permutation test on triangularity (over 1000 permutations) corrected for False Discovery Rate (FDR) (p < 10^−4^). The Principal Convex Hull/Archetypal analysis (PCHA) showed that the triangle was the best polygon to enclose all the data points among planes with 2-6 vertices. In fact, a triangle shape distribution (n=3 vertices) explained the majority of variance (>99.5% variance), and increasing the number of vertices did not improve the amount of variance accounted for (**Figure S1 Supplementary Material**).

**Figure 1.**
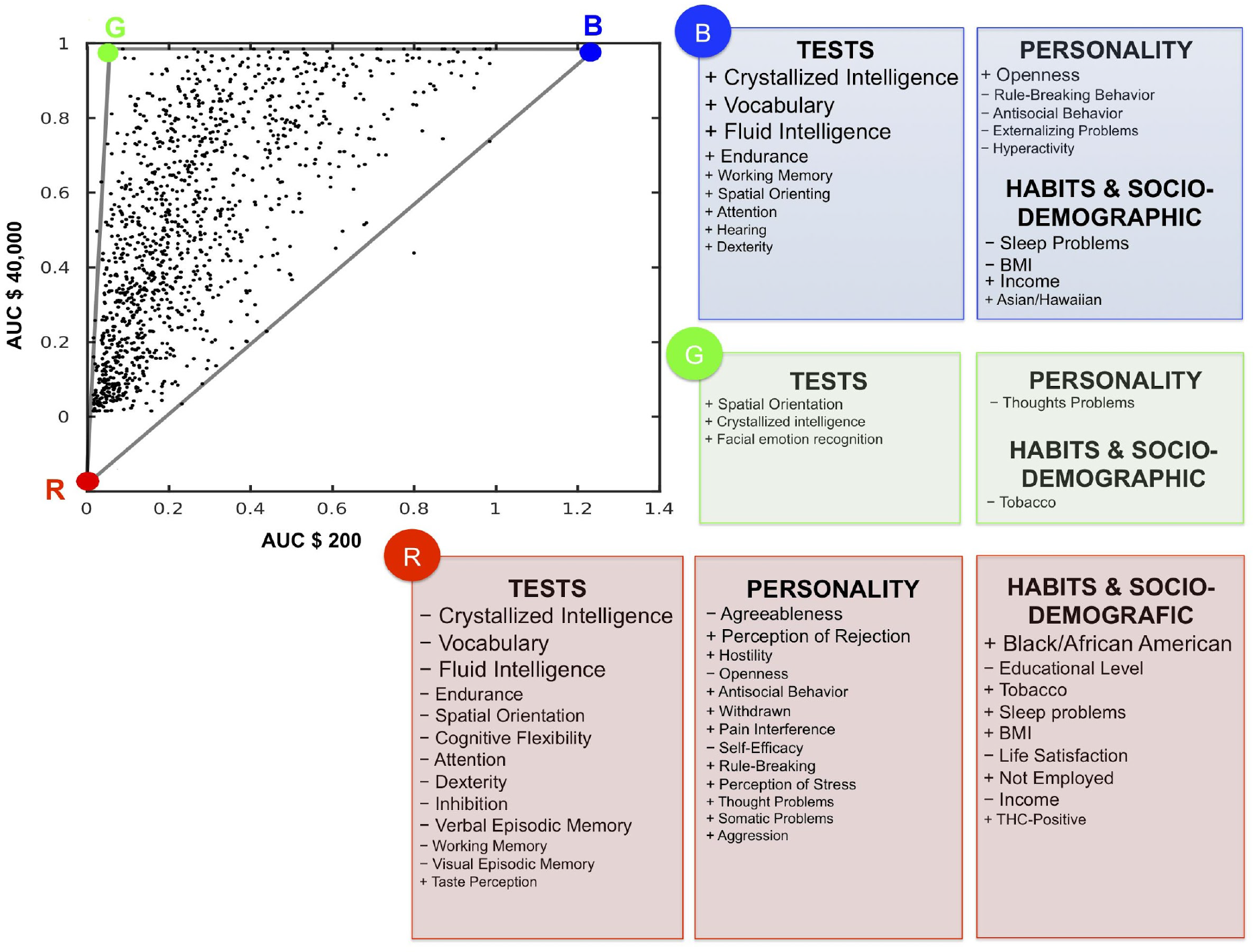
Pareto distribution (triangular polytope) in a space of AUC $200 (x-axis) versus AUC $40,000 (y-axis). The AUCs (Area-Under-the-Curve) are two measures of the Delay Discounting Task. The distribution of AUC scores is triangular hence fitting Pareto optimality theory. The three vertices of the triangle (labeled as Blue, Green, Red) represent the archetypes and indicate three ideal time preferences. Each archetype enriches on specific features that are listed in the adjacent tables. These features include cognitive, sensory and physical abilities, personality traits, habits and socio-demographic variables, and were identified by an enrichment analysis (see also Figure 2 and Table 1). The size of the font corresponds to the relative significance of each trait (larger font, lower p-value).

We further validated the present results performing a split-half replication, in which the analysis was separately run on two independent group of subjects. The only significant triangle that emerged in both groups was that defined by the DDT measures (for both sub-samples: p < 10^−4^, after FDR correction) (**Figure S2**).

Next, we confirmed that this Pareto front distribution was independently significant in subjects of different gender (p <10^−4^ independently for male and female subjects) and race (Asian-Native Hawaiian or Other Pacific populations: p = 5 * 10^−2^; for White subjects p = 10^−4^; for Black or African American individual p = 0.2 (**Figure S3**). In summary, the Pareto front for the DDT is highly significant, and robust over race, gender, and different groups of subjects.

### Archetypes in the Delay Discounting task

The DDT requires an individual to choose between immediate smaller versus delayed larger virtual rewards. Although the rewards are hypothetical, there is a good correspondence of the outcomes with real rewards (Lagorio, & Madden, 2005). Two fixed amounts of money were used as reference for the choice. In half of the conditions the ‘reference’ monetary amount was relatively small ($200), whereas in the other half was high ($40,000). The HCP version of the DDT identifies ‘indifference points’, i.e. where an individual is equally likely to choose a smaller reward earlier (e.g., $50 immediately) versus a larger reward later on (e.g., $200 in 2 years). The AUC discounting measure is a summary measure that provides a valid and reliable index of how steeply people discount delayed rewards (Myerson et al., 2001): smaller is the value, higher is discounting. The AUC measure of the DDT, or time preference for reward, is a reliable indicator of self-control in cases of lower discounting rate (i.e. preference for larger delayed rewards), and impulsive behavior in cases of higher discounting rate (i.e. preference for smaller earlier rewards)(Odum and Baumann, 2010).

The three vertices of the DDT triangle (identified by the colors Blue, Red, and Green, Figure 1) identify *archetypes*, namely subjects who are ‘specialists’, i.e. adopt unique strategies to deal with the discounting task while subjects in the middle of the triangle are ‘generalists’. The Blue archetype corresponds to individuals with stable preference for larger rewards that are delayed in time, independently of the amount. The Red archetype identifies individuals with stable preferences for smaller immediate rewards. The Green archetype includes individuals who prefer delayed rewards only when the amount is very large (i.e., $40,000), whereas prefer taking sooner for smaller amounts ($200).

### Enrichment analysis

The Pareto Front theory predicts that these specialists, who adopt different strategies to solve the DDT task, should show trade-offs in other cognitive tasks or in behavioral traits. To test this prediction, we employed an enrichment or density analysis. This analysis tests for systematic increases or decreases in cognitive or behavioral scores as one moves farther away from different vertices. Statistical significance was assessed by permutation tests in which the labels of the subjects belonging to each archetype were shuffled.

#### Cognitive, Physical and Sensory traits

We carried out the enrichment analysis on 46 features reflecting cognitive, physical, and sensory abilities from 1119 participants, with a complete data set.

We found that near the Blue archetype, several cognitive features enriched including crystallized and fluid intelligence, vocabulary knowledge, working memory, spatial orientation, and attention (Figures 1–2; Table 1; **Figure S4**). For all these measures, individuals close to the Blue archetype showed the highest scores, hence they were superior in these domains. Also measures of sensory and physical abilities enriched near/at the Blue archetype, whose subjects showed the highest levels of hearing function, submaximal cardiovascular endurance, and manual dexterity.

**Table 1.** Enrichment analysis of the archetypes. The first column represents the label of each archetype (B = Blue archetype; G = Green archetype; R = Red archetype). The second and the third columns describe the measure and the corresponding trait enriched, respectively. The resulting p-value is shown in the fourth column and it is specified, in the last column, if the value of each trait is maximum or minimum in the bin close to a given archetype. The asterisk indicates traits that are significantly enriched using a 6-bins analysis.

| Archetype | Experimental Measures | Features | Average difference (p-value) | First bin |
| --- | --- | --- | --- | --- |
| B | ReadEng_Unadj | Crystallized Intelligence | 2.5873E-12 | max |
| B | PicVocab_Unadj | Crystallized Intelligence | 1.1805E-10 | max |
| B | PMAT_ACR | Fluid Intelligence | 2.9223E-10 | max |
| B | NEOFAC_O | Openness | 0.00000285 | max |
| B | PSQI_Score | Sleep problems | 0.00001369 | min |
| B | BMI | Body Mass Index | 0.0000747 | min |
| B | Endurance_Unadj | Endurance | 0.00021863 | max |
| B | SAGA_Income: 8 | High Income | 0.00032978 | max |
| B | ASR_Rule_Raw | Rule-Breaking Behaviour | 0.0012588 | min |
| B | ListSort_Unadj | Working Memory | 0.001287 | max |
| B | Race:Asian/Hawaiian/Oth Pacific | Race | 0.0021611 | max |
| B | DSM_Antis_Pct | Antisocial Behaviour | 0.002465 | min |
| B | ER40_CRT | Emotion Recognition (RTs) | 0.0056397 | max |
| B | SCPT_SPEC | Attention | 0.0063268 | max |
| B | VSPLIT_TC | Spatial Orientation | 0.0077114 | max |
| B | Noise_Comp | Hearing | 0.010801 | max |
| B | Dexterity_Unadj | Dexterity* | 0.010861 | max |
| B | ASR_Extn_Raw | Externalizing | 0.013512 | min |
| B | DSM_Hype_Raw | Hyperactivity | 0.017876 | min |
| B | Taste_Unadj | Taste* | 0.037597 | min |
| G | VSPLIT_TC | Spatial Orientation | 0.0040994 | max |
| G | ASR_Thot_Pct | Thought problems | 0.016071 | min |
| G | Avg_Weeday_Any_Tobacco_7days | Tobacco | 0.017359 | min |
| G | ReadEng_Ageadj | Crystallized Intelligence | 0.031099 | max |
| G | ER40_CRT | Emotion Recognition (RTs)* | 0.13533 | min |
| R | ReadEng_Ageadj | Crystallized Intelligence | 2.5873E-12 | min |
| R | Race: Black/African American | Race | 4.0364E-11 | max |
| R | PicVocab_Unadj | Crystallized Intelligence | 1.1805E-10 | min |
| R | PMAT_ACR | Fluid Intelligence | 2.9223E-10 | min |
| R | Endurance_Unadj | Endurance | 3.8829E-07 | min |
| R | SAGA_Education: 12 | Low Education | 1.0987E-06 | max |
| R | VSPLIT_TC | Spatial Orientation | 2.3749E-06 | min |
| R | SAGA_TB_Still_Smoking | Cigarette Smoking | 7.5111E-06 | max |
| R | Avg_Weeday_Any_Tobacco_7days | Tobacco | 0.00001353 | max |
| R | PSQI_Score | Sleep problems | 0.00007023 | max |
| R | CardSort_Unadj | Cognitive Flexibility | 0.00012633 | min |
| R | SCPT_SPEC | Attention | 0.00021717 | min |
| R | NEOFAC_A | Agreeableness | 0.00029454 | min |
| R | BMI | Body Mass Index | 0.00031656 | max |
| R | LifeSatisf_Unadj | Life Satisfaction | 0.00035019 | min |
| R | Flanker_Unadj | Attention/inhibition | 0.00041816 | min |
| R | SAGA_Employ: 0 | Not Employed | 0.00042451 | max |
| R | SAGA_Income: 1 | Low Income | 0.00077582 | max |
| R | Dexterity_Unadj | Dexterity* | 0.00080057 | min |
| R | PercReject_Unadj | Perception of Rejection | 0.00090289 | max |
| R | IWRT_TOT | Verbal episodic memory | 0.00090433 | min |
| R | THC: True | THC-positive | 0.0010352 | max |
| R | ListSort_Unadj | Working Memory | 0.0012108 | min |
| R | AngHostil_Unadj | Anger Hostility | 0.0026881 | min |
| R | DSM_Anxi_Raw | Anxiety | 0.0031224 | max |
| R | PicSeq_Unadj | Visual episodic memory | 0.0032963 | min |
| R | NEOFAC_O | Openness | 0.0045872 | min |
| R | DSM_Antis_Raw | Antisocial Behaviour | 0.005532 | max |
| R | ASR_Witd_Pct | Withdraw | 0.0064302 | max |
| R | PainInterf_Tscore | Pain interference | 0.0070997 | max |
| R | Selfeff_Unadj | Self-efficacy | 0.0076984 | min |
| R | ASR_Rule_Raw | Rule-Breaking Behaviour | 0.007751 | max |
| R | PercStress_Unadj | Perception of Stress | 0.0078439 | max |
| R | ASR_Thot_Raw | Thought problems | 0.011022 | max |
| R | AngAggr_Unadj | Anger Aggression | 0.011784 | max |
| R | DSM_Somp_Raw | Somatic complains | 0.012219 | max |
| R | ASR_Oth_Raw | Other problems | 0.017609 | max |
| R | Taste_Unadj | Taste* | 0.031563 | max |

**Figure 2.**
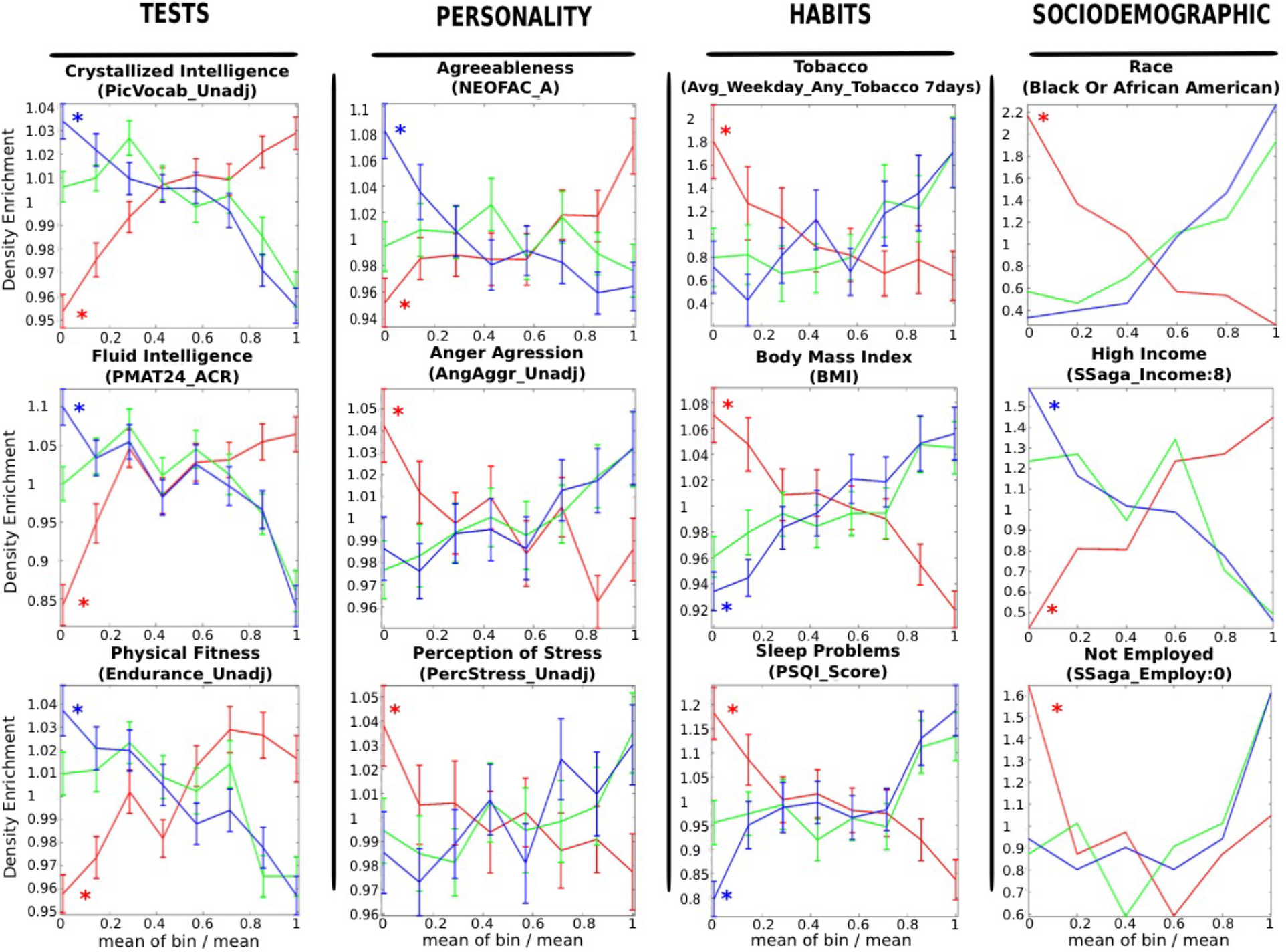
Enrichment of different features near each archetype. Individuals were binned to equal sized bins according to distance from each archetype. The average value in the bin is normalized by the average value in the whole front distribution. The error bars are computed only for continuous measures. The enrichment analysis included cognitive tests, personality scales, habits and sociodemographic features. Curves for features that enrich significantly near an archetype are marked with an asterisk.

When focusing on the Green archetype, individuals near this vertex scored high on measures of cognitive flexibility, crystallized intelligence and spatial orientation, and were the fastest in recognizing facial emotions.

Finally, individuals closest to the Red archetype showed the lowest levels of performance in multiple cognitive domains. These included crystallized and fluid intelligence, vocabulary and spatial orientation, cognitive flexibility, attention and inhibition, working memory, verbal and visual episodic memory. These individuals also manifested the lowest level of performance on endurance and dexterity tasks. However, they showed the highest scores in a taste perception test, i.e. stronger perceived intensity to gustatory stimuli.

Therefore, individuals near the Red archetype showed a lower overall *g* factor. Notably, many of the cognitive, physical, sensory traits (excluding taste perception) reached a minimum near the Red archetype, and increased rapidly with distance from that archetype.

#### Personality, Habits, socio-demographic traits

Data from 1123 participants were analyzed. Two analyses were performed separately on 70 continuous and 40 discrete measures (however, for clarity they will be described jointly).

The enrichment analysis was carried out on measures clustered into: (1) self-reported measures reflecting behavioral, social, and emotional problems, adaptive functioning, and substance use (e.g., ASR and DSM-oriented measures); (2) habits and physiological variables (e.g., quality of sleep, smoking); (3) socio-demographic features (i.e., educational level, race, income) (Figures 1–2; Table 1; **Figure S5**). Individuals closest to the Blue archetype resulted more open to experiences, defined as an appreciation for art, creativity, intellectual curiosity, and a preference for variety and novelty. They also reported the lowest scores on scales related to sleep problems, rule-breaking and antisocial behavior, hyperactivity and externalizing behaviors (such as impulsivity and aggression). Finally, they had the lowest Body Mass Index (BMI), a measure of body fat.

Individuals close to the Green archetype were characterized by minimum scores in thought problems (i.e., hallucinations, strange thoughts and behaviors, obsessive-compulsive behavior, self-harm and suicide attempts), and number of cigarette smoked per day (or other tobacco-related substances (Table 1; **Figure S5**)).

Finally, near the Red archetype, several features enriched with maximum scores in scales reflecting aggressive, hostile, antisocial and rule-breaking behavior, withdrawn behavior and anxiety. Furthermore, individuals closest to the Red archetype reported the lowest life satisfaction, highest perception of stress, most feelings of social rejection, most somatic complaints, most thoughts problems, greatest interference of pain perception in daily life, and poorest sleep quality. Near this archetype, we also observed the highest number of smokers, individuals reporting to smoke the most cigarettes per day, and cannabis users as indicated by the number of positive cases to the THC drug test on the day of the experiment (Figures 1–2). Notably, BMI (obesity) was also maximal in the bin next to the Red archetype, and steeply declined with distance from that archetype.

Examining socio-demographic variables, individuals close to the Blue archetype had the highest income whereas individuals close to the Red archetype had the lowest income, lower educational level, and were most frequently unemployed.

Finally, when considering enrichment on the variable race, Black or African-American individuals were more numerous near the Red archetype, whereas Asian (and Hawaiian or other Pacific Islanders) individuals were more concentrated in the bin closest to the Blue archetype (Figure 2). The variable race was one of the strongest enriched features (p= 4.06×10^−11^). Therefore, it is important to ask whether a triangular distribution for the DDT scores existed separately in each race. As shown above (**Figure S3**), a Pareto optimal distribution was found in each racial group, i.e. when considering separately White, Asian and Hawaiian individuals, or Blacks. In Black subjects, however, the distribution was no longer significant, which is compatible with the results of the enrichment analysis (see **Figure S3**).

In summary, this enrichment analysis shows that stronger (blue archetype) and more flexible (green archetype) self-control, as indexed by the DDT scores, are associated with higher fitness on cognitive, behavioral, socio-economic, and health variables, while weaker self-control is associated with lower scores. Importantly, Blue and Green archetype subjects scored highest on different domains, suggesting different cognitive profiles (Figure 1 and Table 2).

### Structural variables

We examined 56 measures related to mean volume of both white and gray matter, both in specific anatomical brain regions and at the whole brain level, normalized per intracranial volume. Measures were collected from a total of 1105 participants.

The analysis revealed that total cortical gray matter volume was highly enriched near the Blue archetype, reaching a maximum value close to this archetype (Figure 3). In order to compare directly changes in gray matter volume as a function of the archetype, we also ran an ANOVA restricted to individuals close to each of the three vertices (100 participants per group). This analysis showed a significant effect of the archetype [F(2, 297) = 7.9; p < .001; *η*_*p*_^*2*^ = .05], with the Blue archetype being characterized by a larger cortical volume, as compared to both the Red and Green archetypes (*p* < .05; Bonferroni correction) (Figure 3). No difference was instead observed between the Red and Green archetypes (p > .05).

**Figure 3.**
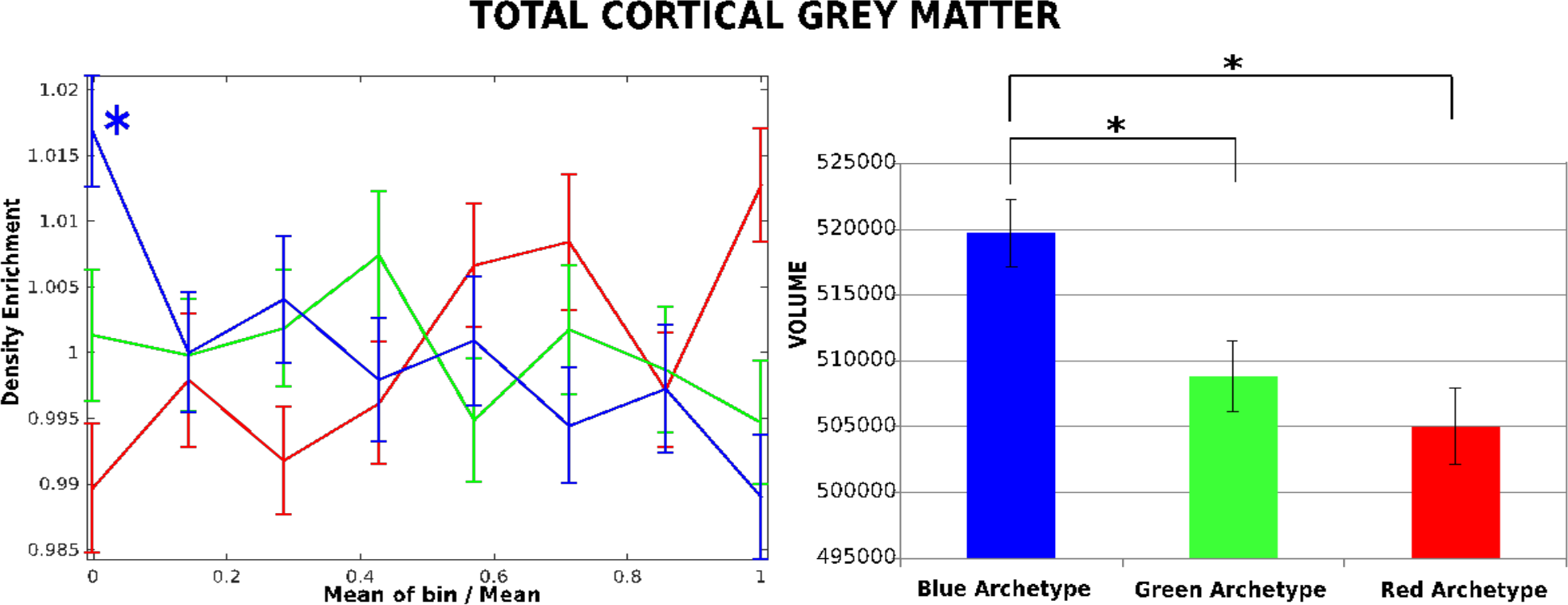
Total cortical grey matter volume varies as a function of archetype. The enrichment analysis (*left panel*) shows that total grey matter volume is enriched for the Blue archetype. The histograms (*right panel*) indicate mean volume in the sub-groups of participants (n=100 for each group) that are closest to the three archetypes. Total cortical gray matter volume is maximal for individuals next to the Blue archetype, intermediate next to the Green archetype, and minimum next to the Red archetype. Asterisks highlight significant differences. Bars indicate standard error.

In summary, stronger self-control (blue archetype) is associated with larger gray matter volume. Importantly, again Blue and Green archetype subjects showed a different profile.

### Brain functional connectivity

In order to explore differences in functional organization we compared resting state FC to/from these regions in three samples of subjects (each n=100) who were closest to each archetype on the DDT. The three samples were matched in gender frequency (percentage of females: Red=63%; Green=52%; and, Blue=57%) (Chi-square test, p > 0.1 for each paired comparison), and age (Average age: Red=28.9 years old; Green=28.6 years old; Blue=29.6 years old) [F(2,299) = 1.99, p > 0.1], variables known to influence functional connectivity.

From the paired hierarchical analysis of connectivity patterns two main clusters emerged: one including regions in medial prefrontal and parietal cortex plus hippocampus, para-hippocampus, and amygdala; the other including basal ganglia, thalamus, and lateral prefrontal cortex (Figure 4A).

**Figure 4.**
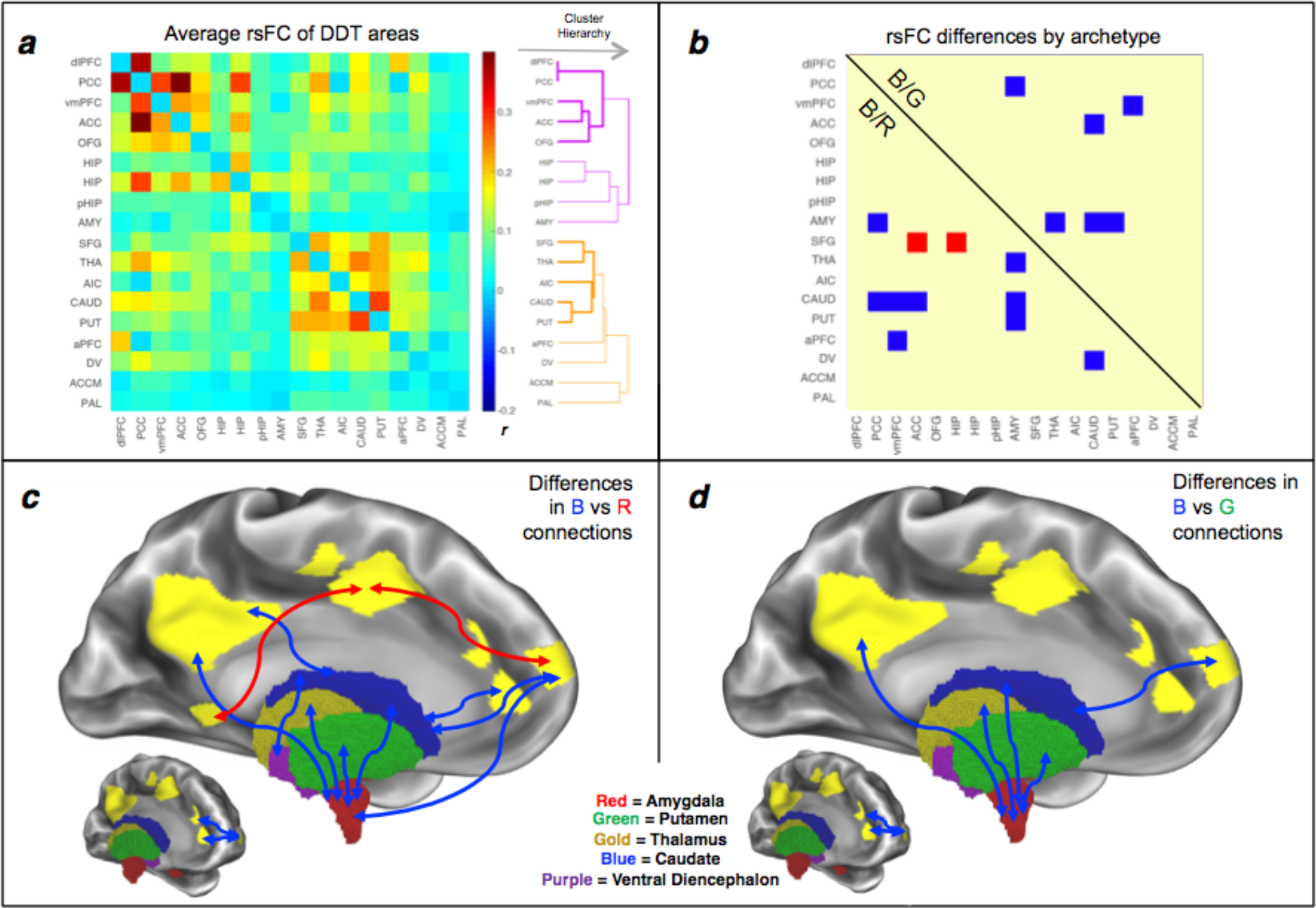
Resting-state functional connectivity differences between archetypes. (***a***) Average rsFC matrix between regions of interest involved in reward and delay discounting task. The FC matrix is divided in two clusters based on a hierarchical cluster analysis (the colour indicates the same functional module membership; the thickness of the line represents the similarity of FC weighted by the connectivity significance). (***b***) Differences in rsFC among the three archetypes as identified by post-hoc comparisons. The lower triangular part compares Blue (B) versus Red (R) archetypes; the upper triangular part contrasted B archetype versus Green (G) archetype. In each part the connections are coloured according to the group with stronger rsFC. (***c*** and ***d***) Spatial representation of connections that differ among archetypes. Significant connections are coloured according to the archetype with higher connectivity level, separately for B/R comparison (c panel) and B/G comparison (d panel). Cortical average position of brain areas involved in archetype-related differences are delineated in yellow while relevant subcortical nuclei are represented with colours specified in the relative legend.

The cortical module (violet in Figure 4A) encompasses areas belonging to the frontoparietal network (FPN) and the default mode network (DMN), typically involved in control-and regulatory-related processes. The subcortical module (orange in Figure 4A) includes regions more strictly related to reward processes.

To examine difference among the three archetypes, we ran a bootstrap-ANOVA. We found mainly FC differences *between* clusters, specifically between prefrontal and cingulate regions, involved in control and regulation, and subcortical regions, involved in reward. In contrast, there were no significant FC group differences in ROI *within* each cluster.

Post-hoc comparisons showed that subjects in the Blue archetype, as compared to subjects in the Red and Green archetypes, had increased FC: *1*) between amygdala and Posterior Cingulate Cortex (PCC), thalamus, caudate nucleus and putamen; *2)* between caudate nucleus and ventromedial Prefrontal Cortex (vmPFC), anterior cingulate cortex (ACC), PCC, amygdala and ventral diencephalic structures (e.g., substantia nigra, hypothalamus, thalamus); and *3)* between anterior Prefrontal Cortex (aPFC) and vmPFC (Figure 4B). All these connections, except those involving the amygdala, were also stronger in subjects of the Green archetype as compared to subjects in the Red archetype. Finally, the Red archetype showed stronger FC between superior frontal gyrus (SFG) and ACC and hippocampus, as compared with the other two archetypes. In summary, stronger (blue archetype) and more flexible (green archetype) self control are associated with stronger FC between reward/emotion related regions (e.g. amygdala, caudate) and control related regions.

### Twin correlations and heritability

Finally, we sought to investigate the heritability of time preferences for rewards by assessing possible differences in intra-class correlations (r) for the AUC $200 and AUC $40000 between pairs of monozygotic twins (MZ; n = 130) and dizygotic twins (DZ; n = 138) by means of Fisher’s z test.

The correlation value did not significantly differ between MZ and DZ pairs, either for the AUC $200 (MZ *r* = 0.30 *versus* DZ *r* = 0.32; *z* = − 0.208 *p* = 0.48), or the AUC $40,000 (MZ *r* = 0.51 *versus* DZ *r* = 0.40; *z* = 1.158 *p* = 0.124).

The difference in MZ-DZ correlation for AUC $ 40,000 was 0.11, indicating a broad heritability (*h*^2^) of only 0.22 (as assessed by Falconer’s formula, see the study by Deary et al., 2009 for a similar approach). For $AUC 200, this calculation was even meaningless as the value for DZ twins was higher than the value for MZ twins. Therefore, MZ twins were not substantially more similar in delay discounting than DZ twins. The heritability (*h*^2^) value indicates that there is not a strong genetic dominance of this trait, as genetic dominance can be inferred for DZ twin correlations that are about ¼ MZ twin correlations.

## Discussion

In the present study we applied Pareto Optimality theory and its methodology to investigate the presence of archetypes in human cognition and behavioral data, which according to the theory reflect evolutionary trade-offs among multiple cognitive and personality traits. Through a data-driven method, we found that, among all the possible combinations of pairs of traits used to project the data points into the morphospace (the space of all phenotypes), one combination stood out over all others, representing the strongest Pareto Front solution. This Pareto Front distribution was obtained when we projected the data derived from two measures of the DDT, a task that measures time preferences for reward, an index of self-control and regulation. This Pareto Front triangular distribution was the only significant to permutation tests on the whole sample of subjects (n=1205), but it was also replicated as the only significant Pareto distribution in smaller samples (in each half of the subjects) (**Figure S2**).

These findings provide important clues to the evolutionary nature of self-control, and time preference for reward in humans in particular. Moreover, they demonstrate that this trait is a crucial dimension to describe the diversity of human behavior and brain structures, offering insights into evolution, cognition, neuroscience, psychology and economy.

### Time preferences for reward: Evolutionary perspective

The evolutionary foundation of time preference for rewards has attracted the interest of economists and biologists for many years (Rogers, 1994). However, this is the first study to demonstrate that time preferences for reward in humans are shaped by natural selection according to the Pareto optimality theory. Three optimally competing strategies, or archetypes, have been identified: The Blue archetype characterized by stable preference for larger rewards that are delayed in time; the Green archetype characterized by preferences for delayed rewards when these are very large; and, the Red archetype characterized by a consistent preference for immediate smaller rewards. The Blue archetype is defined by enriched features that are typically considered positive and desirable qualities, at least in a highly structured and modern environment, thus it is easily understandable the evolutionary advantage for this archetype. For example, being intelligent, agreeable, and open, as well as physically fit, could increase the likelihood to find a mate, as well as earning a high income could increase the offspring quality, via better nourishment and/or investment in education.

Likewise, traits enriched near the Green archetype are advantageous. Green archetype flexibly changes the strategy according to the reward amount, suggesting - respect to the other two archetypes - higher ability in modifying and adapting the behavior on the basis of external environmental features. Also, individuals close to the Green archetype have the most effective emotional expression recognition, thus they are likely to be facilitated in the understanding of others’ feelings and needs.

The evolutionary advantage for the Red archetype appears instead less intuitive at first glance, but it can be accounted for by at least three reasons. The first is the presence of ‘evolutionary mismatch’ between the environment in which we currently live and the environment in which we evolved. Therefore, a behavior that was adaptive and well suited for evolution becomes inappropriate into our current environment (Robson and Samuelson, 2010). In some circumstances, for example, children and adolescents showing aggressive and externalizing behaviors become dominant and respected in their peer groups, whereas in other cases become unpopular or rejected (Frankenhuis and Del Giudice, 2012). Following the same logic, it is conceivable that individually maladaptive traits of the Red archetype are the result of an adaptive strategy developed to achieve social status and dominance.

Second, according to life history theory, time preferences are influenced by resource scarcity, mortality and uncertainty in the environment (Griskevicius et al., 2011). Delay discounting rate was found to be steepest under stressful conditions, in people with low socio-educational background or poor health, all conditions in which individuals close to the Red archetype report to live (Chao et al., 2009; Griskevicius et al., 2011).

Third, natural selection would favor individuals who made reproductive efforts sooner. In this regard, although the HCP dataset does not include such information, we expect that individuals close to the Red archetype are more likely to have their first child sooner and have a larger number of offspring. This suggestion is driven by the evidence that a steeper discounting (evaluated in teenagers and young adults) is associated with a range of sexual behaviors, as having early first experience with sexual intercourse, and past or current pregnancy (Chesson et al., 2006). Furthermore, if discounting rate is influenced by the expected future fitness, then living in relatively adverse circumstances (e.g., elevated risk of mortality, high stress levels, resource scarcity) makes individuals more prone to expend reproductive effort immediately (Daly & Wilson, 2005), as also demonstrated in other species like wasps (Roitberg et al., 1993). Future studies are however needed to explore this relationship.

As for the nature vs. nurture question: are archetypes in time preferences for reward genetically or environmentally determined? We suggest that the absence of significant differences between MZ and DZ correlations and the low heritability (*h*^2^) value indicate a weak genetic influence on this trait, and that environmental influences may come into play. Yet, natural and cultural selection are not mutually exclusive. Heritability of time preferences is indeed not constant across lifespan. It is higher during late childhood/adolescence (Anokhin et al., 2011) and several studies found genetic polymorphisms being associated with differences in time preferences (Boettiger et al., 2007; Eisenberg et al., 2007). By contrast, heritability has less contribution in adulthood (age range of HCP participants: 22-35 years), when other factors, such as environmental stressors and/or cultural factors, could have an impact on individuals’ time preferences to some extent.

In light of this, the high concentration of Black and African American individuals close to the Red archetype might be interpreted as the result of their adverse health and socioeconomic conditions, as consistently revealed by the large amount of data collected through the NSAL (The National Survey of American Life: http://www.rcgd.isr.umich.edu/prba/nsal.htm#overview).

### Cognitive Neuroscience and Psychology Perspectives

Our results do not refuse the concept of g factor: we observed that the Blue archetype has higher abilities in all cognitive domains compared to the Red archetype, which has instead the lowest level of performance. We also found that typically ‘positive’ non-cognitive qualities, as openness, physical fitness, good sleep quality, high income, cooccurred in the Blue archetype, as well as typically ‘negative’ features (aggression, rule-breaking behavior, drug addiction, high BMI, low life satisfaction and self-efficacy, sleep problems etc…) co-occurred in the Red archetype. Therefore, the g factor is not restricted to cognitive abilities, but should be extended also to other behavioral traits of individuals.

Importantly, however, some variables such as cognitive flexibility, recognition of facial emotion, minimal thought disorders, and low smoke use enrich the Green archetype. This indicates that, at least for some variables, trade-offs in time preference shape variability along a third axis beyond a simple g factor positive correlation. Interestingly, in a recent study, Smith et al. (2016) investigated the relationship between individual subjects’ behavioral/cognitive measures in the HCP and functional brain connections in a single analysis. They identified one strong mode of population covariation that can be represented by a “positive-negative” axis. Our study supports and extends Smith et al.’ findings, suggesting that this “positive-negative” axis is linked to self-regulation, and time preferences for reward in particular.

Our study demonstrates that the three archetypes also differ in brain structure and functional connectivity. The Blue archetype has larger cortical gray matter volume respect to the other two archetypes, confirming a possible association of brain volume not only with intelligence (Ritchie et al., 2015), but also with self-control, which is a critical function implied in the DDT (MacLean et al., 2014). Our results extend the findings of a phylogenetic study that has evaluated self-control in 36 species, revealing that the crucial mechanism underlying the evolution of self-control is increases in absolute brain size (MacLean et al., 2014).

Differences among the three archetypes were observed also in the function connectivity profiles of brain regions associated with DDT (Li et al., 2013; Liu et al., 2011; Wesley and Bickel, 2014). The functional organization of these regions clusters in two modules, or systems: cortical prefrontal, cingulate and parietal regions involved in control and regulation, and subcortical regions involved in reward and emotions. Importantly, functional connectivity differences between groups of subjects belonging to different archetypes occurred in the projections that connected different modules.

The present set of results is consistent with a number of dual-system models of decision-making (e.g., Bechara, 2005; Bickel et al., 2007). These models state that decision-making is the result of the relative balance of activation between two neurobiological systems (Bickel et al., 2007). The evolutionarily older impulsive system, which comprises limbic and paralimbic regions (amygdala, ventral pallidum, striatum, nucleus accumbens) values immediate rewards. By contrast, the more recently evolved control system, composed of the PFC and other cortical regions such as ACC, is important for the inhibition/regulation of the impulsive system and the associated valuation of delayed rewards. Our findings support these models, showing that the ability of delaying a reward is associated with a stronger functional coupling between cortical and subcortical regions, which might underlie a more efficient regulatory influence.

More specifically, differences in functional connectivity regarded the connections centered on amygdala, caudate, and aPFC, with subjects in the Blue archetype having stronger connections than subjects in the Red and Green archetypes.

The amygdala is classically considered the core region of emotion regulation (Costafreda et al., 2008), and functions as cognitive-emotional connector hub (Pessoa et al., 2008). In line with our results, altered amygdala-centered connectivity has been found in drug addicts (Sutherland et al., 2012), who frequently show steeper discounting rates and lower self-regulation (Bickel et al., 2011). For example, some studies reported decreased resting-state functional amygdala-centered connectivity in cocaine addicts (Gu et al., 2010), active heroin abusers (Wang et al., 2010), and prescription-opioid addicts (Upadhyay et al., 2010).

Our findings also suggest that the ability to self-control and postpone a reward is the result of a stronger functional connections to/from the caudate nucleus. Fronto-striatal circuitry is indeed implicated in inhibitory control (Ghahremani et al., 2012), and, more specifically, caudate nucleus is associated with behavioral control and goal-directed actions (Grahn et al., 2008). Importantly, Goldstein and Volkow (2011) documented that connection between dorsal caudate and frontal regions facilitates self-control. On the basis of this body of evidence, we propose that caudate nucleus plays a pivotal role in self-control and time preference for reward, possibly acting as a relay structure to connect prefrontal (including ACC) regions for control and regulation, with sub-cortical emotion-and reward-related structures, such as thalamus, amygdala, and substantia nigra, which was recently found to support value coding in the DDT (Zhang et al., 2017).

In contrast, the Red archetype shows stronger functional connections between ACC and superior frontal regions. Although at a first sight this result appears counterintuitive and contradicting the findings discussed above, it is, however, consistent with a recent study that found stronger functional coupling in ACC-frontal circuits to be predictive of a poorer DDT task performance in cocaine users (Camchong et al., 2011).

Finally, from a psychological perspective, although the present study cannot make any conclusion about causal relationships, it provides the most comprehensive overview of the associations between time preference and other individuals’ attributes.

We observed that such trait is able to explain alone individual differences not only in cognitive abilities, but also personality traits, habits and dysfunctional behaviors, sociodemographic features. Notably, a stable preference for immediate smaller rewards seems to predict a constellation of behavioral and real-life problems, including hostile, antisocial, rule-breaking and withdrawn behaviors, anxiety, thought problems, sleep problems, pain and somatic complains, high levels of stress and perception of rejection, low levels of life satisfaction and self-efficacy, high BMI, and substance addiction. Our findings are in line with a large body of findings showing that steeper discounting rates are associated with a range of impulse-control disorders and unhealthy behaviors, drug addiction and lower life satisfaction (Bickel & Mueller, 2009; Odum and Baumann, 2010, for recent reviews). Therefore, time preference appears to be a promising candidate endophenotype for multiple dysfunctional behaviors and might represent a therapeutic target for treating these disease states.

## Acknowledgements

This study was supported by a grant from the University of Padova: PROGETTO STRATEGICO “FC-NEURO” (N°: STPD114NY2) to M.C.

## Contribution

MC, AM, and GC developed the study concept. All authors contributed to the study design. LK and AP performed the data analysis. GC, LK, AM, and MC interpreted the data. GC drafted the article. MC, LK, AB, AP, and AM provided revisions. LK drafted the Supplementary Files. All authors approved the final version of the article for submission.

